# Persistent Memory as an Effective Alternative to Random Access Memory in Metagenome Assembly

**DOI:** 10.1101/2022.04.20.488965

**Authors:** Jingchao Sun, Rob Egan, Harrison Ho, Yue Li, Zhong Wang

## Abstract

The assembly of metagenomes decomposes members of complex microbe communities and allows the characterization of these genomes without laborious cultivation or single-cell metagenomics. Metagenome assembly is a process that is memory intensive and time consuming. Multi-terabyte sequences can become too large to be assembled on a single computer node, and there is no reliable method to predict the memory requirement due to data-specific memory consumption pattern. Currently, out-of-memory (OOM) is one of the most prevalent factors that accounts for metagenome assembly failures. In this study, we explored the possibility of using Persistent Memory (PMem) as a less expensive substitute for dynamic random access memory (DRAM) to reduce OOM and increase the scalability of metagenome assemblers. We evaluated the execution time and memory usage of three popular metagenome assemblers (MetaSPAdes, MEGAHIT, and MetaHipMer2) in datasets up to one terabase. We found that PMem can enable metagenome assemblers on terabyte-sized datasets by partially or fully substituting DRAM at a cost of longer running times. In addition, different assemblers displayed distinct memory/speed trade-offs in the same hardware/software environment. Because PMem was provided directly without any application-specific code modification, these findings are likely to be generalized to other memory-intensive bioinformatics applications.

## 1 BACKGROUND

Ever since the first 454/Solexa sequencers broke the dawn of the next-generation sequencing revolution, the rate of increase in sequencing data has been growing exponentially at a pace exceeding Moore’s law. The number of nucleotide base pairs (bp) in public repositories is estimated to reach the exabase scale (10^18^ bp) before 2025 (Stephens et al., 2015). Metagenomics is one of the main contributors to this rapid growth of data. Metagenomics, the study of microbial genomes directly isolated from their natural habitats, removes researchers from the laborious and time-intensive prerequisite of isolating and cultivating microbes (Tyson et al., 2004; Venter et al., 2004). Powered by next-generation sequencing, metagenomics offers an unprecedented opportunity to gain a deep understanding of the microbial communities around us or within us and to harness their genetic and metabolic potential for our health and environmental safety.

However, the construction of individual microbial genomes from a complex microbial community with thousands of species from billions of short reads faces both data and algorithmic challenges (reviewed in (Ayling et al., 2020)). It was initially thought impossible until pioneering work demonstrated its feasibility (Hess et al., 2011; Iverson et al., 2012). At that time, assemblers developed for single genome assembly were used for metagenome assembly because there were no metagenome-specific assemblers available. Since then, metagenome assemblers have been developed that consider the specific characteristics of metagenomic datasets, such as uneven sequencing depth for different member species. These assemblers include meta-IDBA (Peng et al., 2011), metaSPAdes (Nurk et al., 2017), MEGAHIT (Li et al., 2016), and many others. Several recent studies have provided a comprehensive comparison of the computational performance and accuracy of these assemblers (Sczyrba et al., 2017; Vollmers et al., 2017; Meyer et al., 2021). While most of these assemblers can efficiently take advantage of the modern CPU’s multiple processing capabilities, they are limited on a single computer node and, therefore, are not able to assemble very large datasets due to the limited memory capacity. For terabase-scale metagenome datasets, researchers have very few options. Swapping memory using fast disks or even from multiple machines over a fast network running JumboMem (Pakin and Johnson, 2007) can help if the extra memory required is minimal, but this significantly extends the runtime. meta-RAY (Boisvert et al., 2012) uses MPI to distribute large metagenome assembly to multiple computer nodes. To overcome its limitation that it only assembles very abundant species, hybrid strategies have been developed to first use meta-RAY in a computer cluster to assemble abundant species (which often comprise most of the sequencing data), followed by MEGAHIT or metaSPAdes in a single node to assemble unassembled reads (Wang et al., 2019). Recently, MetaHipMer used UPC++ to assemble very large metagenome datasets with high accuracy and efficiency (Hofmeyr et al., 2020), but it runs best on a supercomputer that is not readily available to most researchers.

New algorithms have the potential to dramatically reduce the memory requirement for metagenome assembly. For example, MEGAHIT uses a data structure called the succinct de Bruijn graph that significantly reduces memory consumption (Li et al., 2016). Since new algorithms take a long time to develop, a more straightforward strategy is to expand the memory capacity of a single system, which does not involve application-specific software development and can be generically applied to other memory-intensive applications. The XSEDE large shared memory system, Blacklight, contains 16TB of shared memory that allows extremely large-scale genome assemblies (Brian Couger et al., 2014). A drawback of this system is that it costs tens of millions of dollars to build. As the price of PC DRAM (DDR4) becomes more affordable, many computer systems are built with several terabytes of DRAM. Intel^®^ Optane™ Persistent Memory (PMem) is a new type of RAM that is packaged in DDR4-compatible modules of 128GB, 256GB, and 512GB capacities, much larger than typical DDR4 modules currently available (16GB, 32GB and 64GB) (Intel, 2020). PMem is three to four times less expensive than DRAM by a factor of three to four folds per GB(MemVerge, 2022), which is promising for memory-intensive applications such as metagenome assembly.

PMem can be configured in one of two modes. “Memory Mode” is volatile and uses the DRAM in the system as cache to improve the performance of the PMem, which has a random access latency approximately four times higher than that of DRAM. An application can use PMem in Memory Mode without code modification. In “AppDirect Mode”, the PMem is nonvolatile, and both DRAM and PMem are addressable as system memory. However, applications must be re-factored to be able to use this mode, which is time-consuming and expensive. MemVerge’s Memory Machine™ is software that runs in the user space on Linux systems to virtualize system memory, so that any application can access PMem in AppDirect mode without code modifications. To make this possible, Memory Machine remaps memory pages so that “hot” data are moved to DRAM and “cold” data are moved to PMem, resulting in overall performance that resembles DRAM. The DRAM capacity reserved for “hot” pages is termed the DRAM tier and can be configured manually in the Memory Machine, or Memory Machine can automatically adjust as the number of “hot” pages changes.

PMem has been successfully applied to many memory-intensive applications, such as databases(MemVerge, 2021; Aerospike, 2019; SAPHANA, 2019; Redis, 2019). This work reports its first application on metagenome assemblies on a server configured with DRAM and PMem. We evaluated the feasibility and performance of the running time and memory consumption of several common metagenome assemblers.

## 2 RESULTS AND DISCUSSION

### 2.1 Detailed memory usage profiling of metagenome assemblers

Modern metagenome assemblers take a “multi-k” approach to assemble species of various abundances in the microbial community. For example, the metaSPAdes pipeline first constructs multiple de Bruijn graphs, each of a distinct kmer size, of all reads using SPAdes. Then it transforms the graphs into the assembly graph, followed by graph simplification and graph traversal to obtain contigs (Nurk et al., 2017). To profile its use of computational resources, we ran metaSPAdes on the 233GB Wastewater dataset and record its CPU, memory. As expected, each phase of the de Bruijn graph construction is computationally and memory intensive, as well as the final assembly graph phase (Figure 1). In any phase where the memory consumption of metaSPAdes is greater than the available DRAM, an Out-of-Memory (OOM) failure occurs. To ensure the success of the pipeline, a workstation is required to have a sufficient memory that is larger than the peak (maximum) memory.

**Figure 1.**
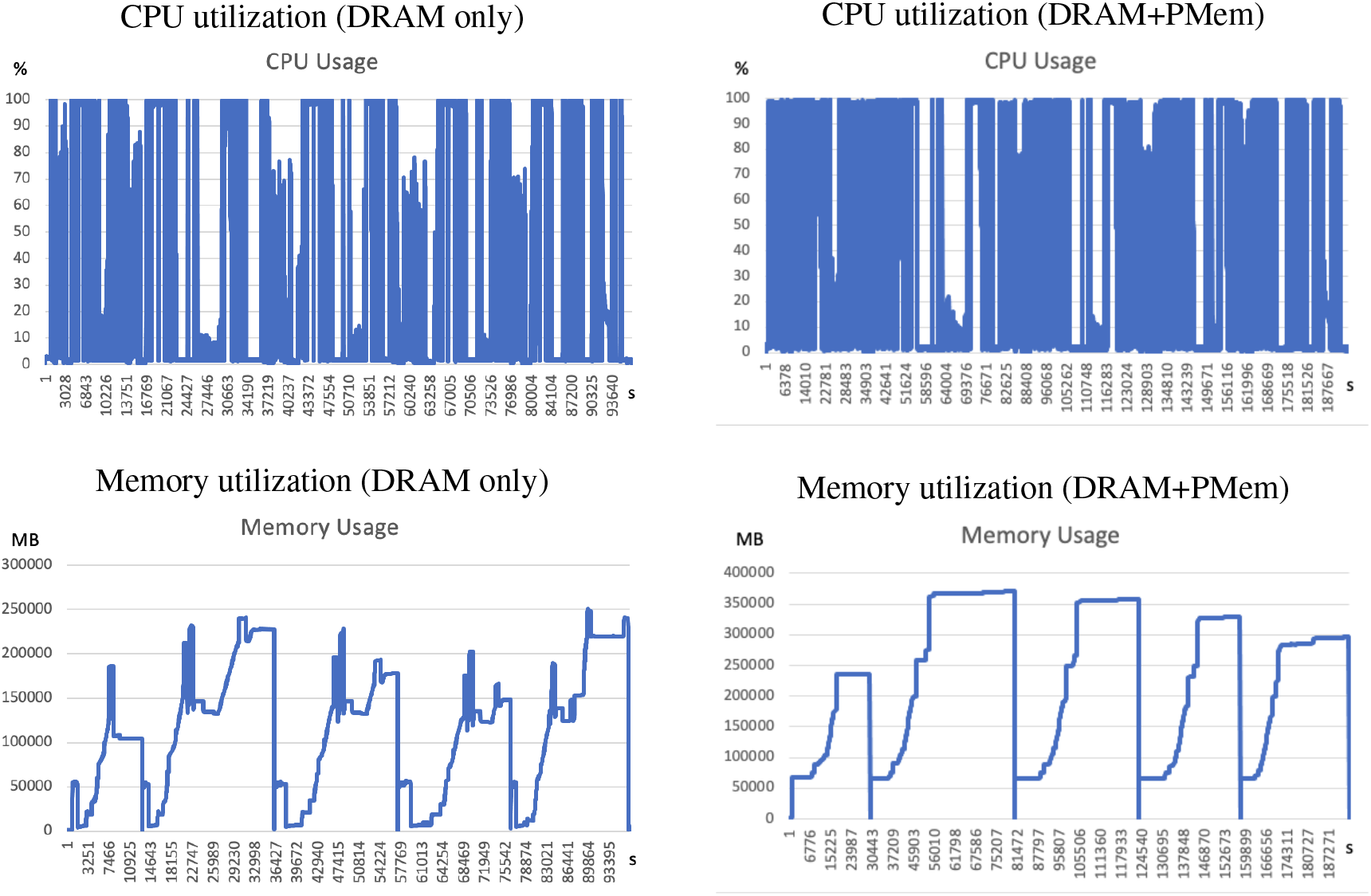
System metrics for metaSPAdes execution on the Wastewater dataset using DRAM-only (left) or using PMem with 32GB DRAM tiering (right). System metrics were recorded for the period that metaSPAdes was running. The CPU utilization timeline (in percentage), memory usage (in MB) are shown. The horizontal axis shows the wall clock time in seconds.

To test whether or not enabling PMem leads to changes in CPU or memory utilization, we used the Memory Machine to configure 32GB of DRAM while supplying the rest with PMem. We saw an increase in peak memory consumption from (250GB to 370GB), which is likely due to the overhead incurred by the Memory Machine software, as the page size was increased from 4KB to 2MB. The general pattern of memory utilization was the same. Furthermore, CPU utilization increases due to additional hot-swapping options (Figure 1).

The memory consumption profiles of MEGAHIT and MetaHipMer2 are similar to metaSPAdes with MEGAHIT requiring the least amount of memory and MetaHipMer2 requiring the most.

### 2.2 PMem can effectively satisfy the requirement of metaSPAdes for large amount of DRAM at a small cost of running speed

The maximum memory consumption to run metaSPAdes on the Wastewater dataset with only DRAM was approximately 250GB, and the run was finished in 26.3 hours. To test whether or not PMem can be used to substitute DRAM for metaSPAdes, we used the Memory Machine to configure decreasing amounts of DRAM and measured PMem usage and running time. We found that PMem can substitute DRAM in all tested memory configurations, resulting in savings in DRAM of up to 100% (Figure 2A). Meanwhile, increasing DRAM savings also led to a longer execution time of metaSPAdes due to the slower performance of PMem compared to DRAM. When 100% PMem is used to replace DRAM, the run time of the metaSPAdes on the Wastewater dataset was approximately twice as long (2.17 ×).

**Figure 2.**
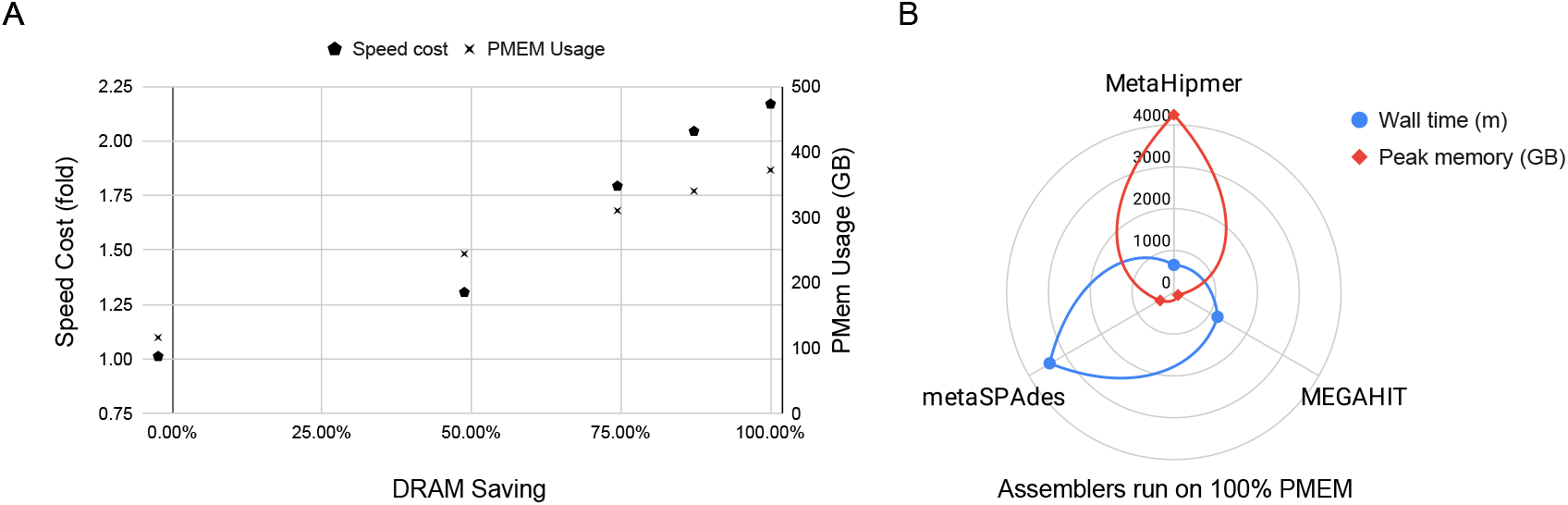
(A) PMem saves DRAM with a speed cost. The percent of DRAM saving is calculated based on the baseline DRAM consumption of running metaSPAdes on the Wastewater dataset (250GB, horizontal axis). The speed cost in folds is shown on the left vertical axis, whereas the amount of PMem used in GB is shown on the right vertical axis. For each memory configuration, we ran the pipeline twice and showed the average results (speed and PMem amounts). **(B)** Comparison of MetaHipMer2, MEGAHIT, and metaSPAdes running on the Wastewater dataset. The three assemblers were run using 100% PMem. The total time of the wall clock (in minutes) is shown in blue, and the maximum memory consumption (in GB) is shown in red.

Running metaSPAdes on the larger 1.2TB Antarctic dataset with only DRAM led to an OOM error, as the required memory exceeds the total amount of DRAM (768GB). When running with Memory Machine with 550GB DRAM tiering, the peak memory reached 1.8TB, and the run was completed successfully in 11.15 days.

### 2.3 PMem supports other metagenome assemblers including MetaHipMer2 and MEGAHIT

In addition to metaSPAdes, we also tested MetaHipMer2 (Hofmeyr et al., 2020) and MEGAHIT (Li et al., 2016) on the Wastewater dataset with 100% PMem (Methods). MetaHipMer2 consumed the most memory (peaked at 4.2TB), while it took the least time (647 min). MEGAHIT used the least memory (peaked at 124GB), but took almost twice as long and finished the assembly in 1,191 min. metaSPAdes used a peak memory of 372GB and took the longest time (3,423 min). The comparison of the three assemblers is shown in Figure 2B.

All three assemblies had roughly the same size and length metrics. About 70% 31-mers are shared among these assemblies. These indicate that all three assembly experiments were successfully completed. Because there is no ground truth reference, no further quality and accuracy evaluations were performed for this experiment.

We did not attempt to compare the three assemblers using the Antarctic Lake Metagenome Dataset because MetaHipMer2 would likely run into OOM. It required 13TB of RAM distributed across multiple nodes in a supercomputer to complete for this dataset in a previous experiment.

## 3 CONCLUSIONS

For terabyte-scale metagenome assembly projects, existing solutions are either expensive (a fat shared-memory machine) or have limited hardware availability (supercomputers). We demonstrated the feasibility of running a large-scale metagenome assembly on commodity hardware by substituting DRAM with persistent memory (PMem). If a running time is not a critical factor, we showed that PMem is a costeffective option to extend the scalability of metagenome assemblers without requiring software refactoring, and this likely applies to similar memory-intensive bioinformatics solutions.

## 4 METHODS

### 4.1 Hardware environment

All experiments were carried out on a single server with 2 Intel(R) Xeon Gold 6248 3.0GHz (Turbo 3.9GHz), each with 24 cores (48 threads). Its memory includes a total of 768GB DDR4 DRAM and 12× 512GB PMem 100 series (6TB total). Six 2TB SSDs were configured as a single 12TB volume. The server was running CentOS 8 Linux with a 4.18.0-305.7.1.el8_4.x86_64 kernel.

### 4.2 Software environment

Memory Machine Release 2.1 by MemVerge Inc. was used to provide memory virtualization. The normal memory page size in the Linux kernel is 4KB. Transparent HugePages (THP) is a memory management system in the Linux kernel that tries to use HugePages (2MB) greater than the default (usually 4 KB). Using large page sizes can improve system performance by reducing the amount of system resources required to access page table entries. Memory Machine implements its own management of HugePages, and thus the default THP was disabled. Memory Machine can be launched by running the command mm before other commands.

### 4.3 Metagenome assemblers

SPAdes (Saint Petersburg Genome Assembler) version 3.15.3 was used. For use with Memory Machine, the source code was compiled using dynamic linking by turning off the options SPADES_STATIC_BUILD and SPADES_USE_MIMALLOC in the make file. Furthermore, the OpenMP scheduling was changed from dynamic to static (Line 255 in /src/common/utils/kmer mph/kmer index builder.hpp). The following metaSPAdes command line options were used “-only-assembler”, “-k 33,55,77,99,127”, “-meta”, “-t 96”.

MetaHipMer2 (MHM2) version 2.1.0.37-g01c2b65 was used. The source code for MHM2 was compiled with UPC++[upcxx-ipdps19] version 2021.3.0, which in turn was built using the included ‘install_upcxx.sh’ script that builds UPC++ using the default ‘smp’ conduit for Symmetric Multi-Processor and shared-memory for communication between processes. MHM2 was executed using the default options on the dataset: ‘mhm2.py-r filtered_wgs_fastq.fastq’

MEGAHIT version v1.2.9 was used. The source code was downloaded and compiled following the instructions on (https://github.com/voutcn/megahit).

### 4.4 Datasets

The following data sets were used:

1. The Wastewater Metagenome Dataset was downloaded from NCBI (BioProject: PRJNA506462, Accession: SRR8239393). It was derived from samples taken at a waste water treatment plant in Idaho (Stalder et al., 2019). This dataset has a total uncompressed size of 164.8GB.
2. The Antarctic Lake Metagenome Dataset was downloaded from the JGI GOLD database with the accession no Gs0118069 (https://gold.jgi.doe.gov/study?id=Gs0118069). It was derived from samples taken from two meromictic lakes in Antartica (Wang et al., 2019). This dataset has twelve files totalling 1.37TB.

## 5 ACKNOWLEDGMENTS

The authors thank Brian Foster for his tips on running metaSPAdes. Rob Egan and Zhong Wang’s work is supported by the US Department of Energy Joint Genome Institute (https://ror.org/04xm1d337), a DOE Office of Science User Facility, supported by the Office of Science of the US Department of Energy operated under Contract No. DE-AC02-05CH11231. The Antarctic Lake Metagenome Dataset was generated from the JGI CSP proposal DOI: 10.46936/10.25585/60000788. Harrison Ho’s work is supported by a fellowship from the National Science Foundation Research Training Program (NRT) in Intelligent Adaptive Systems.

